# Advanced Melissopalynology For Ensuring Honey Authenticity And Quality: Insights From Diverse Floral Resources as Diverse Natural Component to Apitherapy

**DOI:** 10.1101/2025.07.13.664608

**Authors:** Aysan Torkamani, Melika Radnejad, Mahla Pirzadeh, Homa Zarabizadeh, Narjes Rezazadeh moghaddam, Helia Hajihassani, Ahmadreza Mehrabian

## Abstract

As the global honey market experiences exponential growth, ensuring the authenticity and quality of honey has become a critical concern. This study employs melissopalynology, the analysis of pollen in honey, to ascertain both the botanical and geographical origins of honey samples. Through the examination of honey samples representing a diverse range of plant families, the research highlights the prevalence of *Asteraceae* and *Fabaceae*. The findings emphasize the pivotal role of melissopalynology in safeguarding honey quality, facilitating fair pricing, and detecting adulteration. This research not only enhances the authentication cycle for monofloral honey but also contributes significantly to the creation of a comprehensive melissopalynological database, underpinning future studies in honey authenticity and quality control.

## 1. Introduction

The development of the honey market through the world identification of botanical origin, as well as its authenticity, has received much attention in international society (Kaškonienė and Venskutonis, 2010). Fake and low-quality honey are significant issues in the global honey trade. Authenticity assessment of honey is extremely important for consumers.

Melissopalynology, as an efficient method as well as a primary standard, has been extensively used to determine the purity, geographical, and botanical origins of natural honey (Cornalba and Griffiths, 1975, Alvarez-Suarez, 2017). Founded this method for the first time. Although this method has been associated with shortcomings. Despite its shortcomings, various laboratory procedures have since been developed to study pollen grains in honey (Bryant Jr and Jones, 2001). At the meetings of a working group organized at the International Honey Commission of Apimondia, which established some changes to reduce the variability and increase the accuracy of the method (Von Der Ohe et al., 2004). Harmonized methods of melissopalynology have been developed in the form of scientific papers.

Generally, honey is classified as monofloral (mainly originating from a single plant species) or polyfloral (produced from multiple plant species). (Belay et al., 2017). In addition, the botanical origin of honey influences the physical, chemical, sensory, and bioactive features of honey (Kortesniemi et al., 2018) That is considered a valuable indicator of the quality and authenticity of honey. Up to now, several studies have been done on melissopalynology around the world, following Southwest Nigeria (Ebenezer and Olugbenga, 2010, Adeonipekun, 2012, Agwu et al., 2013), Bulgaria (Atanassova et al., 2012), Lithuania (Čeksterytė et al., 2013), Argentina (Forcone et al., 2005, Estevinho et al., 2012, Costa et al., 2013), Turkey (Erdoğan and Erdoğan, 2014, Ozler, 2015), Mexico (Villanueva-Gutiérrez et al., 2009), Spain (Pardillo López and Serna Ramos, 2007), Brazil (Oliveira et al., 2010), Malaysia (Rosdi et al., 2016), Oman (Sajwani et al., 2007), Algerian (Nair et al., 2013), Australia (Sniderman et al., 2018), Poland (Stawiarz and Wróblewska, 2010), Ethiopia (Bareke and Addi, 2019), Iran (Shakoori and Salmanpour, 2024).

The climatic diversity and ecological isolation (Frey and Probst, 1986), complicated geological history (Stöcklin, 1968), diverse and unique soils (Hedge and Wendelbo, 1978) The connection of phytogeographical zones (Takhtajan, 1986, Zahran and McKee, 2010) Make Iran a global diversity center for plants (Barthlott et al., 1996, Kier et al., 2005, Davis, 1994) Iran includes about 9500 plant taxa, 30 percent of which are limited to the geographical boundaries of the country (Mehrabian et al., 2015, Mehrabian et al., 2020). Accordingly, Iran has a wide range of high-quality rangelands that contain valuable nectar-producing and medicinal plants, which are considered potential sources of high-quality honey. With a production of 113,000 tons of honey per year, Iran is ranked as the world’s third-largest honey-producing country. Also, about 55 different types of honey are produced in Iran. To date, approximately 54 types of monofloral honey have been identified in Iran. *Ziziphus spina-christi, Astragalus* spp., *Thymus* spp., *Eryngium* spp., *Echinops* spp., *Medicago sativa, Citrus* spp., *Tillia begonifolia*, and *Eucalyptus calamadonensis* are the primary sources of monofloral honey in Iran.

To achieve comprehensive honey quality control, the authenticity of the honey, determination of its herbal origin, health evaluation, and organoleptic evaluation are all integrated, and each part of the honey quality control puzzle is completed. Melissopalynology, as one of the essential criteria for determining honey authenticity and its botanical origin, plays a vital role in controlling the quality of honey, ensuring fair pricing, identifying fraud, and highlighting the medicinal properties of honey. In addition, The botanical origin greatly influences the of chemical compounds in honey, which in turn influences the medicinal properties of honey. Therefore, determining the botanical origin as well quality control is greatly influential in the therapeutic properties of honey and apitherapy.

Currently, there is little attention given to melissopalynology and monofloral honey in Iran. It is essential to enhance the processes for authentication and quality control, as well as to establish a grading system for honey pollen. This study aims to: (1) enhance the authentication process for monofloral honey in Iran, (2) analyze the palynological spectra of various Iranian monofloral honey types, and (3) support the development of a comprehensive melissopalynological database in Iran.

## 2. Materials and Methods

### 2.1. Network Analysis

To create this diagram, we utilized VOSviewer software. The data were sourced from ScienceDirect and were specifically chosen to emphasize the connections between melissopalynology, monofloral honey, and polyfloral honey, all within the context of assessing honey quality and authenticity. The diagram reflects research spanning from 2015 to 2025. At the core of the honey map lies the central theme of our study, which focuses on exploring honey quality and authenticity through the lens of melissopalynology. This network map illustrates the interrelationships between authenticity, the active compounds in honey, phenolic compounds, chemometrics, honey authentication, melissopanology, food authentication, physicochemical properties, and other relevant factors. By analyzing the created networks, we can evaluate the proximity and connections between these topics. For instance, in the context of verifying honey authenticity, the networks suggest that this area encompasses the study of fraud detection and food authentication (**Figure 1a**). Upon examining this image, it becomes evident that the topics explored in recent years, such as chemical markers, chemical control, pollen identification, authentication, sensory properties, and related areas, are highlighted in yellow. In contrast, the shift toward darker shades of green signifies research themes that have been the focus of earlier studies (**Figure 1b**). The information presented highlights the significance and the necessary steps required to initiate the targeted project effectively.

**Fig. 1.**
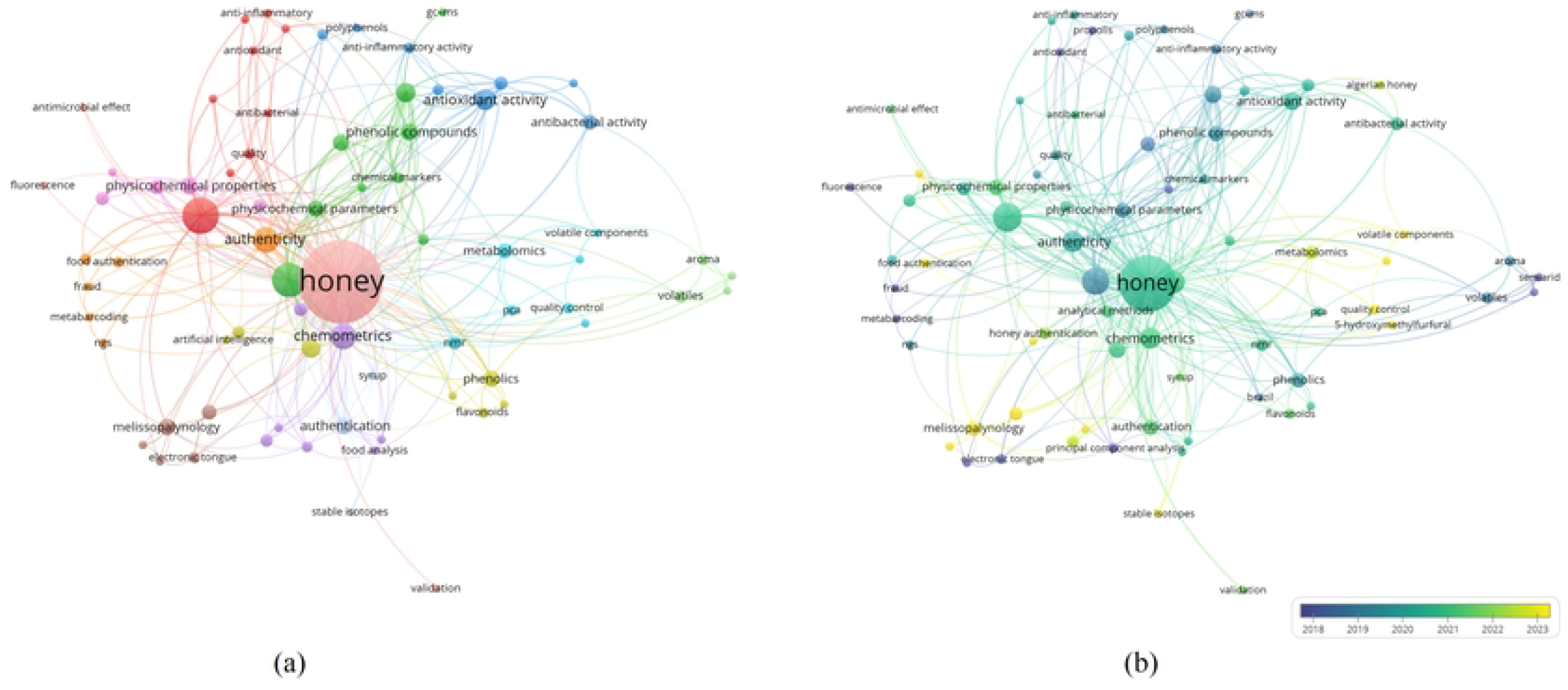

### 2.2. Study Area

The studied area is situated within Iran’s geographical borders, encompassing a total surface area of 1.6 million km^2^ at latitudes ranging from 25° to 40° N and longitudes from 44° to 64° E. Iran is a mountainous country, including some prominent topographic mountain chains such as Alborz, Zagros, and Kopet Dagh, as well as some interior mountain chains. The Alborz topographic zone system is made of a gentle sinus with an east-west orientation in northern Iran that shapes a topographic barrier to the central Iranian plateau and the Caspian Sea (Stöcklin, 1974). The Zagros, as another prominent topographic zone, makes a mountainous ecosystem with a northwest (Eastern Turkey) to the southeast (Makran Mountains) direction (Homke, 2007). The Kopet-Dagh is situated at the eastern boundaries of the Caspian Sea, which continues into northeastern Iran, northern Afghanistan, and Turkmenistan (Buryakovsky et al., 2001). The natural geographical barriers that surround Iran prevent moisture from entering the country’s interior, resulting in a heterogeneous rainfall pattern across Iran. Iran is mainly located in the Mediterranean (western, northwestern Iran, temperate (northern Iran), and tropical (southern coast zones of the Persian Gulf and the Gulf of Oman) macro-bioclimates (Rivas-Martínez et al., 1999). Also, hyper-arid (35.5%), arid (29.2%), and semi-arid (20.1%) climatic zones are affected. In addition, the average rainfall in Iran (860mm) is about one-third of the average rainfall in the world (Amiri and Eslamian, 2010). Iran mainly covers the Irano-Turanian region, Sudano-Zambian, and Euro-Siberian phytogeographic areas of the world, respectively (Takhtajan, 1986) (**Figure 2**).

**Fig. 2.**
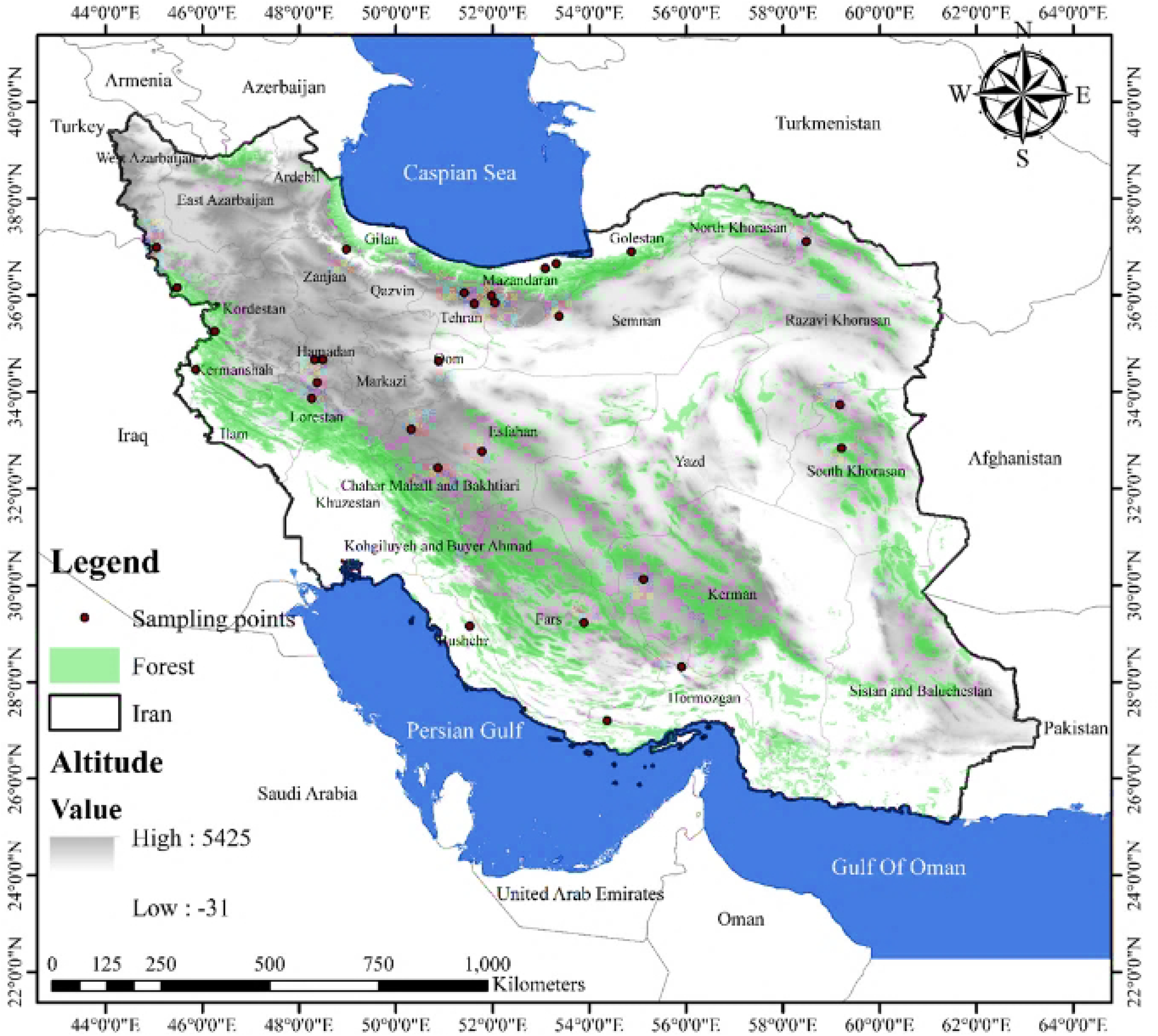

### 2.3. Sample collection

The honey samples were obtained from various apiaries in different geographical regions of Iran. In each sample, 100 g of honey was stored in dark-sealed containers. 42 honey samples were obtained during 2022-2023. The extraction procedure follows the Louveaux method (Louveaux et al., 1978). Accordingly, 10 g of honey was dissolved in 10 mL of warm distilled water, and 50 mL of ethyl alcohol was added to the solution. The solution was centrifuged at 3000 rpm for 10 min. Afterward, the supernatant was rejected, and the remaining sediment solution was diluted again with 10 ml of distilled water and centrifuged again. This process was repeated three times to remove as much sugar and other particles as possible from the Pollen sediment. The pollen grains were transferred to a Neobar slide, and at least 500 pollen grains per sample were used to determine the general number of pollen grains as well as frequency classes per sample (Moar, 1985). Accordingly, the frequency class includes predominant pollen (>45%), secondary pollen (16%–45%), crucial minor pollen (3%–15%), minor pollen (0%-3%), and absent pollen (<1%). The dried pollen samples were mounted on stubs using double-sided adhesive tape and coated with gold using a desktop DC Magnetron sputter coater. The samples were then photographed using an SEM (Cam Scan Hitachi SU3500) **(Figure 3)**. To accurately identify pollen grains, reference materials such as the PalDat website and additional literature sources were consulted

**Fig. 3.**
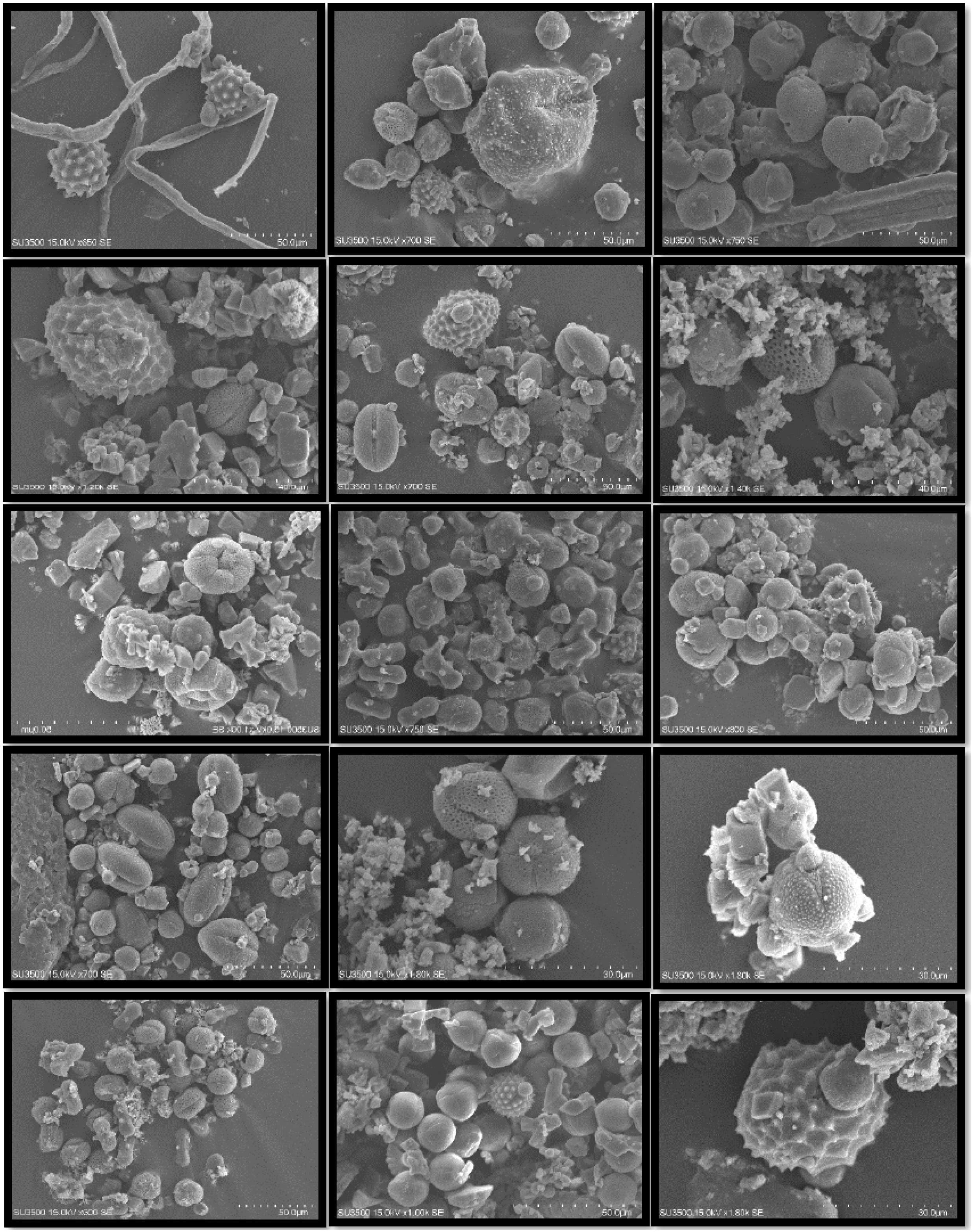

## 3. Results

A total of 23 plant families were identified from the 49 honey samples analyzed using the melissopalynology method. The most abundant families included Apiaceae (25.36%), Fabaceae (19.98%), Berberidaceae (18.63%), and Asteraceae (12.82%). Tilioideae contributed 6.46%, while Brassicaceae and Alismataceae represented 5.76% and 5.71%, respectively.

Several families exhibited moderate to low abundance, such as Boraginaceae (0.75%), Caprifoliaceae (0.75%), Myrtaceae (0.97%), and Plantaginaceae (0.32%). Minor contributions were recorded for families including Solanaceae (0.38%), Rosaceae (0.32%), Liliaceae (0.22%), Onagraceae (0.54%), and Lamiaceae (0.65%).

The least abundant families, each contributing less than 0.1%, were Araceae, Calophyllaceae, Loranthaceae, Malvaceae, Moraceae, Poaceae, and Primulaceae. These findings highlight the diversity of floral resources utilized by honeybees, with significant contributions from key families supporting nectar and pollen availability (**Figure 4**).

**Fig. 4.**
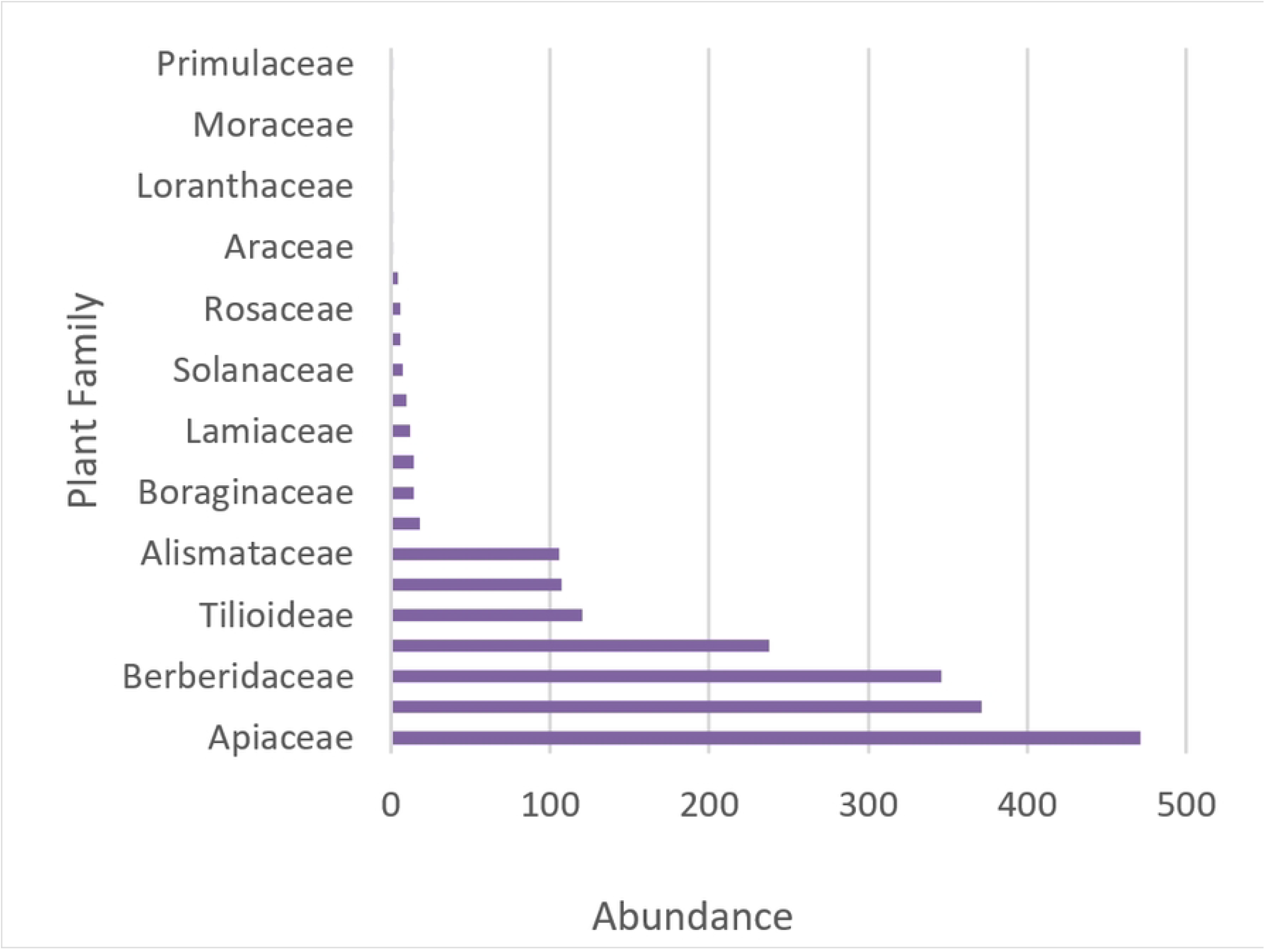

The analysis of 49 honey samples revealed a varied representation of plant families based on their occurrence across the samples. Asteraceae was the most frequently identified family, appearing in 29 samples, followed by Fabaceae (23 samples), Brassicaceae (17 samples), and Apiaceae (16 samples). These families likely represent the primary floral resources utilized by the honeybees in the studied area.

Moderate occurrences were noted for the Boraginaceae family, which appeared in seven samples, along with Alismataceae and Berberidaceae, each represented in six samples, and Myrtaceae, found in five samples. Families such as Caprifoliaceae, Lamiaceae, Plantaginaceae, and Rosaceae were observed in four samples each, while Onagraceae was noted in three samples. Less frequently recorded families included Liliaceae and Tilioideae, each appearing in two samples, alongside Solanaceae, which also had two samples.

Several families were rarely detected, appearing only once in the dataset. These included *Araceae*, Calophyllaceae, Loranthaceae, Malvaceae, Moraceae, Poaceae, and Primulaceae, highlighting their limited contribution to the floral composition of the honey samples.

This distribution underscores the diversity of floral resources in the environment and honeybees’ reliance on a few dominant families for nectar and pollen (**Figure 5**).

**Fig. 5.**
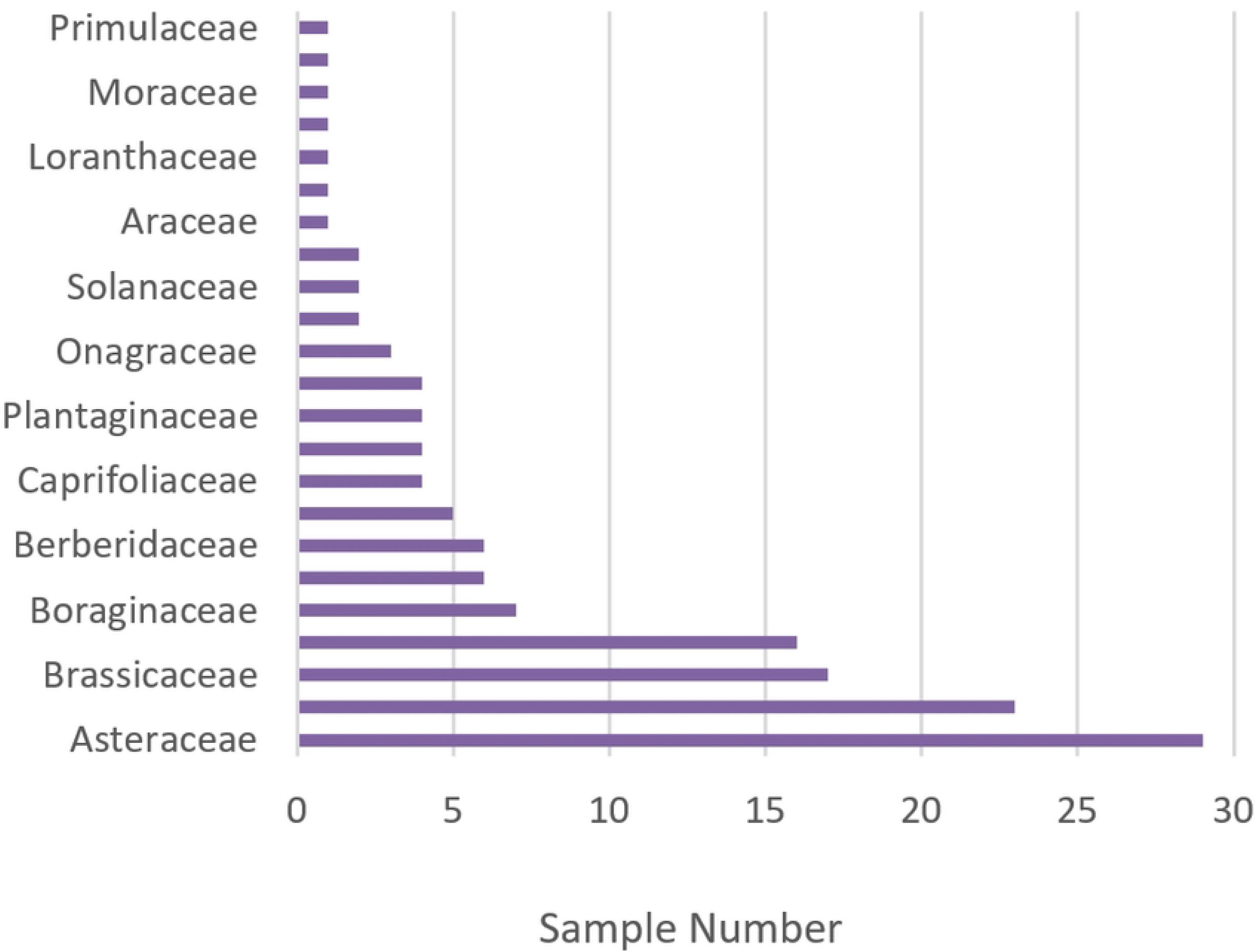

Among the plant families analyzed, a few exhibited the strongest correlations, indicating a significant degree of co-occurrence. Myrtaceae and Alismataceae demonstrated the highest correlation (r = 0.878, p < 0.001), suggesting shared ecological factors or overlapping floral resources. Similarly, Caprifoliaceae and Boraginaceae (r = 0.805, p < 0.001) displayed a strong relationship, likely reflecting complementary contributions to honey composition.

Liliaceae and Berberidaceae (r = 0.712, p < 0.001) also showed a high correlation, potentially indicating their combined availability in specific habitats. Moderate correlations were observed between Rosaceae and families such as Calophyllaceae, Loranthaceae, and Moraceae (r = 0.623 for all, p < 0.001), further highlighting their significant interdependence. These results highlight the considerable influence of specific floral resources on shaping the overall honey composition, driven by their shared environmental and pollinator interactions (**Table 1**).

**Table 1.** Pairwise Pearson Correlations in Plant Families rea.

| Sample 1 | Sample 2 | N | Correlation | 95% CI for $\rho$ | P-Value |
| --- | --- | --- | --- | --- | --- |
| Myrtaceae | Alismataceae | 49 | 0.878 | (0.792, 0.930) | 0.000 |
| Asteraceae | Apiaceae | 49 | 0.257 | (-0.026, 0.502) | 0.075 |
| Brassicaceae | Apiaceae | 49 | 0.761 | (0.611, 0.859) | 0.000 |
| Fabaceae | Apiaceae | 49 | 0.499 | (0.253, 0.684) | 0.000 |
| Rosaceae | Araceae | 49 | 0.623 | (0.415, 0.770) | 0.000 |
| Boraginaceae | Asteraceae | 49 | 0.297 | (0.018, 0.534) | 0.038 |
| Brassicaceae | Asteraceae | 49 | 0.284 | (0.003, 0.523) | 0.048 |
| Caprifoliaceae | Asteraceae | 49 | 0.539 | (0.304, 0.712) | 0.000 |
| Lamiaceae | Asteraceae | 49 | 0.407 | (0.142, 0.617) | 0.004 |
| Myrtaceae | Asteraceae | 49 | 0.279 | (-0.002, 0.520) | 0.052 |
| Liliaceae | Berberidaceae | 49 | 0.712 | (0.538, 0.827) | 0.000 |
| Caprifoliaceae | Boraginaceae | 49 | 0.805 | (0.678, 0.886) | 0.000 |
| Plantaginaceae | Boraginaceae | 49 | 0.660 | (0.465, 0.794) | 0.000 |
| Fabaceae | Brassicaceae | 49 | 0.365 | (0.093, 0.586) | 0.010 |
| Plantaginaceae | Calophyllaceae | 49 | 0.264 | (-0.018, 0.508) | 0.067 |
| Rosaceae | Calophyllaceae | 49 | 0.623 | (0.415, 0.770) | 0.000 |
| Lamiaceae | Caprifoliaceae | 49 | 0.363 | (0.091, 0.584) | 0.010 |
| Plantaginaceae | Caprifoliaceae | 49 | 0.677 | (0.489, 0.805) | 0.000 |
| Malvaceae | Lamiaceae | 49 | 0.686 | (0.501, 0.811) | 0.000 |
| Plantaginaceae | Loranthaceae | 49 | 0.264 | (-0.018, 0.508) | 0.067 |
| Rosaceae | Loranthaceae | 49 | 0.623 | (0.415, 0.770) | 0.000 |
| Rosaceae | Moraceae | 49 | 0.623 | (0.415, 0.770) | 0.000 |
| Onagraceae | Myrtaceae | 49 | 0.355 | (0.082, 0.579) | 0.012 |
| Plantaginaceae | Onagraceae | 49 | 0.401 | (0.136, 0.613) | 0.004 |
| Primulaceae | Plantaginaceae | 49 | 0.264 | (-0.018, 0.508) | 0.067 |
| Rosaceae | Primulaceae | 49 | 0.623 | (0.415, 0.770) | 0.000 |
| Tilioideae | Rosaceae | 49 | 0.290 | (0.009, 0.528) | 0.043 |

The map illustrates the spatial distribution of plant family abundance across the studied region in Hasselt. The gradient is designed to represent the distribution of plant families across different regions visually. Blue signifies areas with the highest diversity, featuring up to eight plant families, while red indicates regions with the lowest diversity, comprising only one plant family. This variation likely reflects environmental and ecological factors, such as soil composition, climate, and vegetation diversity, that influence the availability and diversity of floral resources. Regions with higher family numbers (blue) suggest richer biodiversity and more favorable conditions for diverse plant growth. In contrast, a s with lower family numbers (red) might indicate harsher environmental conditions, limited floral diversity, or dominance of a few plant families. This information is crucial for understanding the ecological dynamics of the region and for designing conservation strategies to maintain or enhance biodiversity (**Figure 6**).

**Fig. 6.**
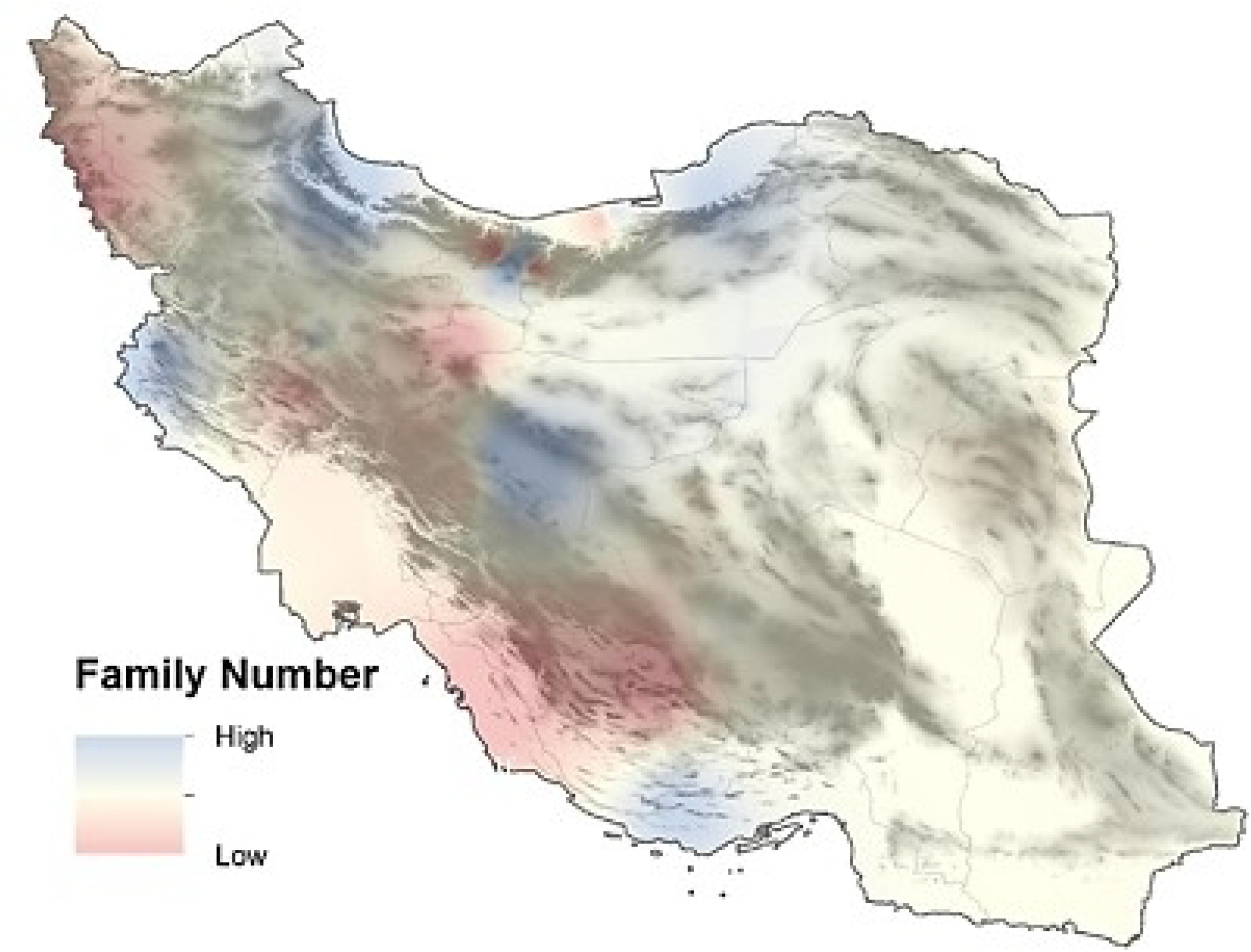

Among the 49 honey samples analyzed, 15 samples were classified as monofloral, with more than 50% of their pollen content belonging to a single plant family. Six plant families were identified as contributing to the monofloral composition of the honey: Apiaceae (three samples), Fabaceae (three samples), Asteraceae (five samples), Berberidaceae (two samples), Brassicaceae (one sample), and Alismataceae (1 sample). Asteraceae was the most frequently associated family with monofloral honey, reflecting its widespread availability and high pollen production. Geospatial analysis revealed that most monofloral honey samples were collected from regions within or near the Elburz and Zagros mountain ranges. These areas are characterized by rich floral diversity and favorable climatic conditions, which provide a habitat for plant families that significantly contribute to honey production. The spatial distribution of monofloral honey samples highlights the ecological importance of mountainous regions in sustaining diverse floral resources and supporting apiculture activities **(Figure 7)**.

**Fig. 7.**
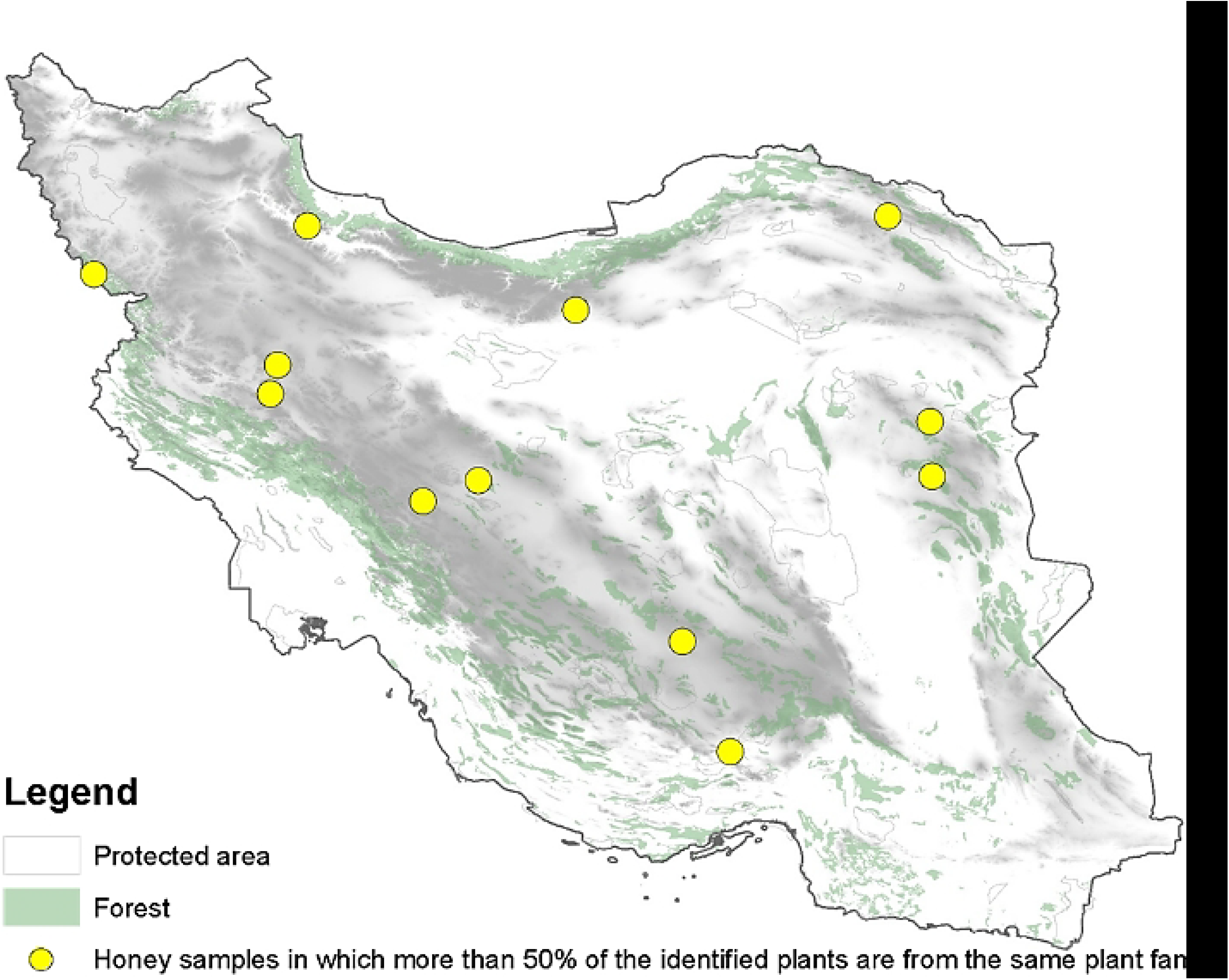

## 4. Discussion

We considered conducting this study to facilitate the recognition of honey and pollen grain origins using the melissopalynology method, which has been shown to achieve 50% efficiency with the aid of the PalDat database. We utilized melissopalynology to identify pollen sources, inform environmental studies, and enhance the quality control of honey, among other applications. The analysis of 49 honey samples revealed a varied representation of plant families based on their occurrence across the samples. The study’s results indicated that a total of 23 plant families were identified from the 49 honey samples analyzed using this method. Apiaceae and Fabaceae were the most abundant families (45.34%), while Araceae, Calophyllaceae, Loranthaceae, Malvaceae, Moraceae, Poaceae, and Primulaceae were the least abundant (less than 0.1%) from the mentioned digits; the diversity and availability of pollen and floral resources are concluded. Correlation analysis revealed strong co-occurrence patterns, particularly between Myrtaceae and Alismataceae (r = 0.878, p < 0.001), suggesting potential ecological interactions or overlapping floral resources. Almost 30.6% of the honey analyzed in this study was monofloral, with the majority of the monofloral honey sources originating from the Asteraceae family, indicating high pollen production. Our results show that the most abundant plant family was Apiaceae, and Asteraceae was the most frequently identified family across the samples. On the other hand, Fabaceae has been assigned the same grading, both in terms of its abundance in the studied area and honey samples as a pollen source. Therefore, we can conclude that there is a direct connection between the most abundant plant family and the family found in the samples the most, although not necessarily a direct one.

In comparison with previous studies, our results align with global melissopalynological studies that emphasize the dominance of Fabaceae, Asteraceae, and Apiaceae in honey composition. Similar trends have been observed in honey from Turkey (Erdoğan and Erdoğan, 2014), Mexico (Villanueva-Gutiérrez et al., 2009), Spain (Pardillo López and Serna Ramos, 2007), and Brazil (Oliveira et al., 2010). However, the unique geographical and climatic conditions of Iran, with its diverse climatic regions ranging from Mediterranean and temperate zones to hyper-arid and semi-arid areas, contribute to a distinct floral composition, particularly with the presence of Berberidaceae and Alismataceae, which are less frequently reported in international studies. Compared to Iranian studies (Shakoori and Salmanpour, 2024). Our dataset offers a more detailed classification, underscoring the importance of region-specific pollen databases. Iran’s diverse climatic regions, ranging from Mediterranean and temperate zones to hyper-arid and semi-arid areas, play a crucial role in shaping the composition of honey. The high presence of the Apiaceae and Asteraceae families is attributed to their widespread distribution in Iran’s montane and steppe ecosystems, which serve as key sources of nectar. The predominance of monofloral honey samples in the Alborz and Zagros mountain ranges underscores the ecological importance of these regions for beekeeping. (Alborz and Zagros are hotspots of Iran’s biodiversity. Zagros habitats 3642 plant species, 792 genera, and 122 native plant families, whereas Alborz is a habitat of 3617 plant species in 876 genera and 146 native plant families) (Shakoori and Salmanpour, 2024). The low occurrence of families like Rosaceae, Moraceae, and Poaceae indicates limited nectar availability or selective foraging behavior by bees.

The aims of the study, as mentioned before, were to improve the authentication cycle in monofloral honey of Iran, analyze the palynological spectra of some Iranian monofloral honey, and promote the melissopalynological database in Iran

For future research, our findings suggest that enhancing honey quality in Iran requires the development of pollen reference databases. Future studies should also incorporate molecular techniques alongside melissopalynology for enhanced species identification at the species level. By establishing a comprehensive classification system for Iranian honey, this study contributes to ensuring fair market pricing, preventing fraud, and highlighting the medicinal properties of high-quality honey varieties. This study plays an important role in completing the apitherapy chain by screening, quality control, and determining the botanical origin of Iranian honeys.

## Abbreviations

SEM: Scanning Electron Microscope
DC: Direct Current
RPM: Revolutions Per Minute
km^2^: square kilometers
r: correlation coefficient
CI: Confidence Interval
N: sample size
ρ: population correlation coefficient
ml: milliliters
g: grams
p: probability value

## Acknowledgments

The authors extend their gratitude to Shahid Beheshti University for its support and contributions to this research.

